# Immune profiling of gliomas reveals a connection with Tau function and the tumor vasculature

**DOI:** 10.1101/2020.07.17.208165

**Authors:** Teresa Cejalvo, Ricardo Gargini, Berta Segura-Collar, Pablo Mata-Martínez, Beatriz Herranz, Diana Cantero, Yolanda Ruano, Daniel García-Pérez, Ángel Pérez-Núñez, Ana Ramos, Aurelio Hernández-Laín, María Cruz Martín-Soberón, Pilar Sánchez-Gómez, Juan M. Sepúlveda-Sánchez

**Author notes:** Equal contribution (Teresa Cejalvo, Ricardo Gargini). Lead Contacts, Dr. Juan-Manuel Sepúlveda-Sánchez, Neurooncology Unit, Hospital Universitario 12 de Octubre, Av Andalucía S/N 28041, Madrid, Spain., Dr. Pilar Sánchez-Gómez, Neurooncology Unit, Instituto de Salud Carlos III-UFIEC, Crtra/Majadahonda-Pozuelo, Km 2, Majadahonda, 28220, Madrid, Spain. Authorship: Experimental design: PSG, JMS, TC, RG; Patient selection, clinical procedures and informed consent process: JMS, DGP, APN, AR, MCMS; Carried out the experiments or molecular analyses: TC, BSC, PMM, BH, DGP, DC, YR, AH; Experiment supervision: PSG and RG; Data analysis: TC, RG, PSG, JMS; All authors have been involved in the writing of the manuscript, and have read and approved the final version.

## Abstract

**Background:** Gliomas remain refractory to all attempted treatments, including those using immune checkpoint inhibitors. The characterization of the tumor (immune) microenvironment has been recognized as an important challenge to get a mechanistic explanation for this lack of response and to improve the therapy of glial tumors.

**Methods:** We designed a prospective analysis of the immune cells of gliomas by flow cytometry. Tumors with or without *isocytrate dehydrogenase 1/2* (*IDH1/2*) mutations were included in the study. The genetic profile and the presence of different molecular and cellular features of the gliomas were analyzed in parallel. The findings were validated in mouse glioma models.

**Results:** We observed that few immune cells infiltrate mutant *IDH1/2* gliomas and we distinguished two different profiles in their *IDH1/2* wild-type counterparts. The first one has an important immune component, particularly enriched in myeloid cells with immunosuppressive features. The second group is more similar to mutant *IDH1/2* gliomas, with few immune cells and a less immunosuppressive profile. Notably, we observed a direct correlation between the immune content and the presence of vascular alterations, which were associated with a reduced expression of Tau, a microtubule-binding-protein that controls the formation of tumor vessels in gliomas. Furthermore, overexpression of Tau was able to reduce the immune content in orthotopic mouse glioma models, delaying tumor growth.

**Conclusions:** There is a correlation between vascular alterations and the immune profile of gliomas, which could be exploited for the design of more successful clinical trials with immunomodulatory molecules.

**Key points:**

1. Mutant *IDH1/2* gliomas harbor few immune cells in the tumor microenviroment.
2. We distinguished two different profiles in the *IDH1/2* wild-type gliomas.
3. There is a correlation between Tau expression, vascular alterations and the immune profile.

**Importance of the Study:** In the present work we have confirmed that IDHmut gliomas are “cold” tumors and we have identified a subgroup of IDHwt GBMs that also contains a low immune infiltrate. By contrast, a large subgroup of IDHwt GBMs are characterized by an important immune component, particularly enriched in myeloid cells, and an elevated expression of the ligand of PD-L1 in the immune compartment. Notably, we have observed a direct correlation between the immune content and the presence of vascular alterations, as well as with the reduced expression of Tau, a microtubule-binding protein that we described as a negative regulator of angiogenesis. Here, we add that overexpression of Tau reduces the immune content in orthotopic glioma models, delaying tumor growth.

This correlation between the vascular phenotype and the entrance and/or the function of the immune cells on gliomas, where Tau could play a central role, opens new venues to find synergistic treatments.

## Introduction

Gliomas are classified as lower-grade gliomas (LGGs) (grade II and III) or grade IV gliomas (glioblastomas (GBMs)). Among the first, detection of 1p/19q co-deletions discriminates oligodendrogliomas from astrocytomas, the latter being enriched in *α-thalassemia/mental retardation syndrome X-linked* (*ATRX*) mutations. By contrast, all GBMs have an astrocytic lineage and they accumulate diverse genetic alterations. Gliomas must be now classified based on the presence or absence of *isocitrate dehydrogenase 1/2* (*IDH1/2*) mutations, as this is a strong prognosis indicator^1^.

In the last decade, immunotherapy with checkpoint inhibitors (ICIs) has been remarkably successful across several tumor types. By contrast, recent clinical trials using anti-programed cell death 1 (PD-1) antibodies in recurrent GBM has shown very few responses^2^, even though the ICIs seems to reach the brain^3^. Compared to responsive cancers, gliomas harbor a lower burden of somatic mutations, fewer infiltrative T cells and a more immunosuppressive tumor microenvironment (TME)^4-6^. These factors could explain why glial tumors remain largely refractory to ICIs. Nonetheless, gliomas are far from being a homogenous entity and disparities in their immune content might also condition their response to different therapeutic strategies. In order to classify the immunological profile of glial tumors, several groups have used computational and immunohistochemical analysis approaches^7-10^. Here, we have performed a comprehensive characterization of the tumors by flow cytometry. Our group have recently described that Tau, a microtubule-binding-protein, impairs the neovascularization of gliomas^11^ and here we have assessed if overexpression of Tau additionally reduces the immune content in the human samples and in orthotopic glioma models, modifying tumor growth.

## Materials and Methods

### Patient samples

All patients gave written informed consent and the study was performed with the approval of the Ethical Committee at “Hospital 12 de Octubre” (CEI_14/023 and CEI_18/024).

### Molecular characterization of the tumors

We used a custom Ampliseq (PCR-based) gene-targeted NGS panel that analyze 30 genes that were previously demonstrated to be frequently mutated in gliomas^12^. DNA from formalin-fixed paraffin-embedded (FFPE) tumor tissues was extracted using the QIAamp DNA FFPE Tissue Kit (Qiagen). DNA was quantified using a Qubit2.0 Fluorometer (Thermo Fisher). Libraries were constructed from 10ng of DNA using the Ion-AmpliSeq Library-Kit v2.0 (Thermo Fisher), according to the manufacturer’s instructions. Libraries were multiplexed, submitted to emulsion PCR and loaded into the chip using the Ion Chef System. Libraries were sequenced using Ion GeneStudio S5 system (Thermo Fisher) according to the manufacturer’s instructions, at average target panel coverage of 800X.

### MGMT Methylation-Specific PCR

O6-methylguanine-DNA-methyltransferase (MGMT) promoter methylation analysis was perfomed as previously reported^12^.

### Flow cytometry analysis

Tumors suspensions were obtained after mechanical and enzymatic disaggregation (Accumax (Merck Millipore) (15 min, room temperature (RT) and filtered through 70μM nylon mesh cell strainer (Fisher Scientific). Erythrocytes were lysed with Quicklysis buffer (Cytognos) and cells were incubated with hFcR Blocking (Miltenyi), previous to antibody (Supplementary Table 2) incubation (20 min at 4°C in PBS 1% fetal bovine serum (FBS)). Viable cells were labelled with a Fixable Viability Stain (Becton Dickinson) (20 min, RT). The analysis was conducted in a Macsquant10 flow Cytometry (Miltenyi). Subset definition was: Neutrophils: CD45^+^CD11b^+^CD16^+^CD15^+^CD14^-^CD33^-^; MDSCs: CD45^+^CD11b^+^CD16^+^CD15^-^CD14^+/-^ CD33^+^; Macrophages: CD45^+^CD11b^+^CD16^-^CD15^-^CD14^+/-^CD33^-^MHCII^+^; Tregs: CD45^+^CD3^+^CD4^+^CD25^+^CD127^lo^.

### Western Blot

The protein extract was generated by mechanical disaggregation using lysis buffer (50 mM Tris-pH 7.5, 300 mM NaCl, 0.5% SDS, and 1% Triton X-100) (15 min with agitation at 95°C). Protein content was quantified using BCA Protein Assay Kit (Thermo-Fisher-Scientific) and 20 μg of proteins were resolved by 10% or 12% SDS-PAGE and then transferred to a nitrocellulose membrane (Amersham Biosciences). The membranes were blocked (1h, RT in PBS and 0.1% Tween-20 with 5% skimmed milk) and then with the primary (Supplementary Table 2) (overnight 4°C) and the secondary (HRP-conjugated anti-mouse or -rabbit, DAKO) (2h, RT) antibodies diluted in PBS-T. Proteins were visualized with ECL (Bio-RAD) using the Imager 680 (Amersham).

### Immunohistochemistry (IHC)

Samples were fixed in 10% formalin overnight, dehydrated through a series of graded ethanol baths and then infiltrated with paraffin. 5 μm thick sections were obtained in a microtome and then sections were rehydrated and permeabilized (1% triton X-100). Antigen retrieval was performed with Citrate Buffer (10mM pH 6) in a pressure cooker (2 min). Endogenous peroxidase inhibition and blocking with normal horse serum was also performed before the incubation with primary antibodies (Supplementary Table 2) (overnight, 4°C) and biotinylated secondary antibodies (15 min). Sections were then incubated with SAV-HRP (10 min) and with DAB (3 min).

### IHC quantification

The IHC score was judged from 0 (no staining) to 4 (the strongest positive staining) in 10 high magnification pictures from each sample. For the quantification of the vasculature we counted the number of dilated vessels per high-magnification field and the relative area covered by the CD34 positive staining. For the latter, 6 fields per sample were counted using the ImageJ program and applying the vascular density plugin.

### qRT-PCR assay

RNA was extracted from the tissue using RNA isolation Kit (Roche). Total RNA (1μg) was reverse transcribed with PrimeScript RT Reagent Kit (Takara). Quantitative real-time PCR was performed using the Light Cycler 1.5 (Roche) with the SYBR Premix Ex Taq (Takara) and specific primers for each gene (Supplementary Table 3). Gene expression was quantified by the delta-delta Ct method. IDH1/2 were evaluated by IHC and PCR.

### Cell culture

The GL261 murine glioma cells were maintained in DMEM plus 10% FBS, 2mM L-glutamine, 0.1% penicillin (100 U/ml) and streptomycin (100 μg/ml).

### Mouse model study

Animal experiments were reviewed and approved by the Research Ethics and Animal Welfare Committee at “Instituto de Salud Carlos III” (PROEX 244/14 and 02/16), in agreement with the European Union and national directives. Intracranial transplantation to establish orthotopic allo-grafts was performed injecting 50.000 cells (resuspended in 2 μl of culture medium) with a Hamilton syringe into the striatum of C57Bl/6 mice (A–P, −0.5 mm; M–L, +2 mm, D–V, −3 mm; related to Bregma) using a Stoelting Stereotaxic device.

### In-silico studies

For studies of gene expression and gene profiling, the merge TCGA dataset (LGG+GBM) was used with 1153 enrolled patients and a set of 702 patients with LGG and GBM tumors with RNAseq values (IlluminaHiSeq). The different immune cell’s signatures (Activated CD8 cell, Central memory CD4 cell, Central memory CD8 cell, Regulatory cell, Type 1 helper cell, Type 2 helper cell, Macrophages, MDSC and Neutrophils) were obtained from 13. For the GSEA (Gene Set Enrichment Analysis) study, the Tau/MAPT expression was continuously computed through the LGG+GBM data-set using the expression by RNAseq, then we used the method continuous class label and genesets from the “CGP: chemical and genetic perturbations” and “CP: BioCarta gene sets”. For correlation studies, the expression values were obtained from http://gliovis.bioinfo.cnio.es. For gene ontology analysis, the DAVID gene ontology 6.8 program was used with a cluster of 500 genes.

### Statistical analysis

The quantified data were represented as mean ± SEM, compared between two groups using the two-tailed Student’s t-test. Differences are presented with statistical significance or p-value (*p <0.05; **p <0.01; ***p <0.001 and ns, not significant). For the correlation analysis between each protein we used the Pearson’s correlation coefficient (R2); P-values were calculated using the GraphPad program. For survival analysis, we used the Kaplan-Meier method and log-rank test using the SPSS program.

## Results

### Identification of different subgroups of gliomas based on the immune profile

Samples from 28 patients diagnosed with glioma (Table 1) were dissociated and analyzed by flow cytometry (Individualized data in Supplementary Table 1). Tumors were classified based on the histology (GBM vs LGG) and based on the presence of IDH mutations. As expected, the majority of LGG were IDHmut (7/9), whereas only 3 out of 19 GBMs carried these mutations. The gating strategy described in Supplementary Fig. 1 was used to characterize different immune populations. In agreement with the literature^14^, there was a significant increase in the number of CD45+ cells in IDHwt GBMs compared to IDHmut GBMs or to LGGs (Fig. 1A). This increase was observed in both the lymphoid (Fig. 1B) and the myeloid (Fig. 1C) components. Among the IDHwt GBMs we identified a subgroup of tumors with a low content of CD45+ cells (less than 15% of the cellular suspension) (herein called GBMwt_lo) (Fig. 1D), with a very similar percentage of immune cells to the one measured in IDHmut tumors (either LGGs or GBMs) and very different from the other group of IDHwt GBMs (herein called GBMwt_hi), where CD45+ cells account for almost 50% of the tumor content (Fig. 1E). Similar differences between the defined glioma subtypes were obtained when we measured independently lymphoid (Fig. 1F) or myeloid (Fig. 1G) cells.

**Table 1.** Characteristics of the study population.

|  |  |
| --- | --- |
| **N** | **28** |
| **Age (years)**<br>Median<br>Range | 52 years<br>30- 82 years |
| **Gender**<br>Female<br>Male | N= 12; 43%<br>N= 16; 57% |
| **Grade of resection**<br>Complete<br>Partial | N= 23; 82%<br>N= 5; 18% |
| **KPS after surgery**<br>100<br>90<br>80<br>≤ 70 | N= 10; 36%<br>N= 9; 32%<br>N= 4; 14%<br>N= 5; 18% |
| **Histological Diagnosis**<br>Astrocitoma<br>Oligodendroglioma | N= 25; 89%<br>N= 3; 11% |
| **Tumor grade**<br>II<br>III<br>IV | N= 3; 11%<br>N= 6; 22%<br>N= 19; 67% |
| **IDH 1/2**<br>Mutated w/t | N= 10; 36%<br>N= 18; 64% |
| **ATRX**<br>Mutated<br>w/t | N= 8; 28%<br>N= 21; 72% |
| **MGMT**<br>Methylated<br>Unmethylated<br>Unknown | N= 21; 72%<br>N= 2; 7%<br>N= 5; 21% |
| **TERT promoter**<br>Mutated<br>Wild Type<br>Unknown | N= 19; 62%<br>N= 10; 34%<br>N= 1; 4% |
| **1<sup>st</sup> line therapy**<br>Stupp (RT + temozolomide)<br>Temozolomide<br>RT + PCV<br>None | N= 21; 75%<br>N= 1; 4%<br>N= 2; 7%<br>N= 4; 14% |

PCV: procarbazine, lomustine, vincristine.

**Figure 1.**
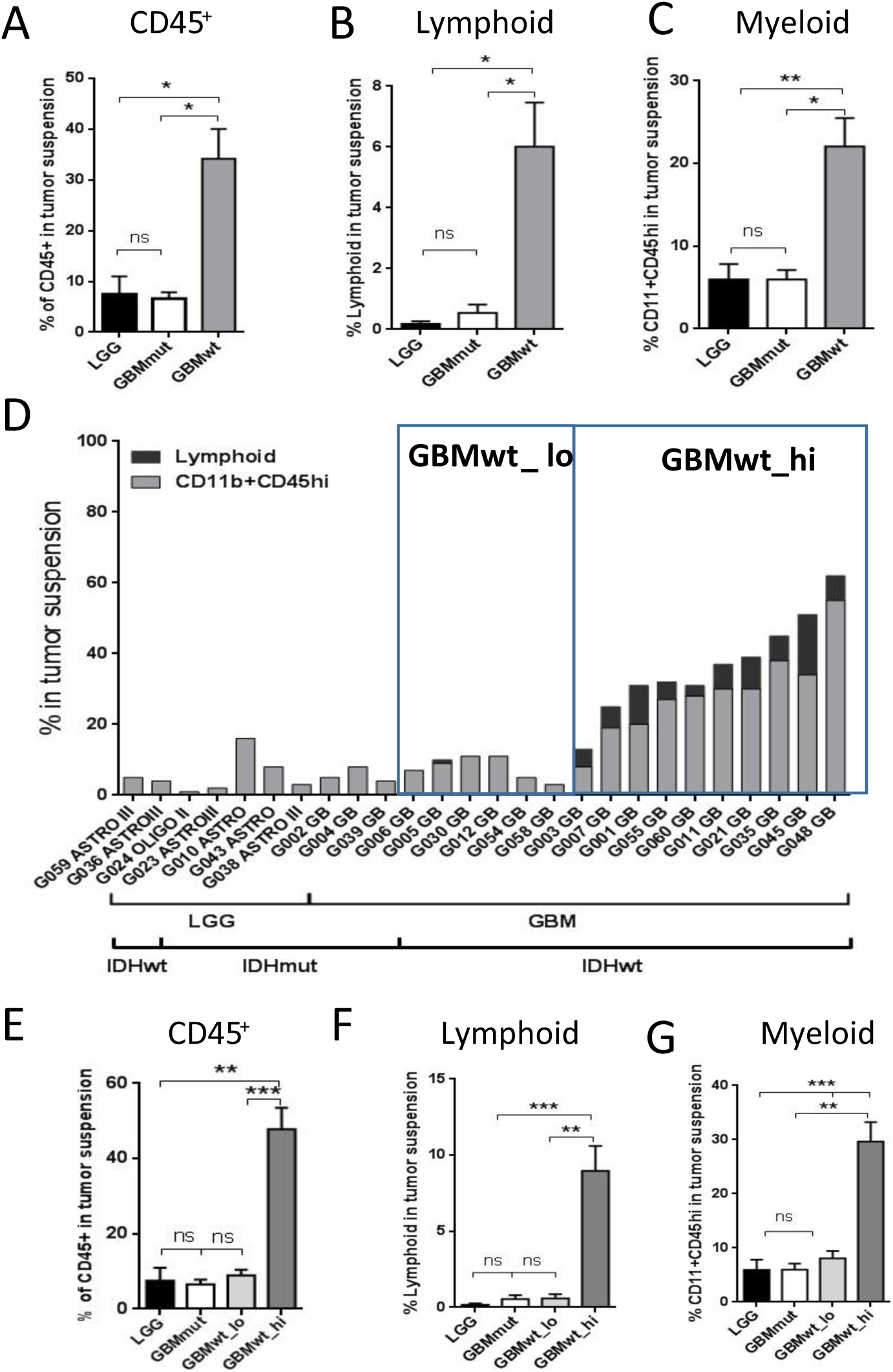
Flow cytometry analysis of the immune infiltrate of gliomas. (A-C) Percentage of CD45^+^ (A), lymphoid (CD45^+^CD11b^-^SSC^lo^) (B) and myeloid (CD45^+^CD11b^+^SSC^lo^ or SCC^hi^) (C) cells on total tumor suspensions in LGG (n=6) and GBM expressing mutant (GBMmut) (n=3) and wild-type (GBMwt) (n=17) IDH. (D) Percentage of lymphoid or CD11b^+^CD45^hi^ populations on total tumor suspension in individual samples. (E-G) Percentage of CD45^+^ cells (E), lymphoid (CD45^+^CD11b^-^SSC^lo^) (F) and myeloid (CD45^+^CD11b^+^SSC^lo^ or SCC^hi^) (G) cells on total tumor suspensions in the four groups of gliomas: LGG (n=6), GBMmut (n=3), GBMwt_lo (n=6), GBMwt_hi (n=11). * p<0.05, **p<0.01, ***p<0.001, ns: not significant.

We performed an analysis of the genetic profile of the tumors. As expected, *EGFR* alterationss were enriched in IDH1wt GBMs, whereas *TP53* mutations were common among the IDH1mut gliomas (Supplementary Fig. 2A). However, we did not detect clear differences between the genetic profile of GBMwt_lo and GBMwt_hi tumors. Regarding the clinical data, patients carrying IDH mutations survived longer than the wild-type counterparts. However, there was no significant differences in the clinical behavior of patients from both GBMwt immune subgroups (Supplementary Fig. 2B).

### Characterization of the myeloid and the lymphoid compartments in the different subgroups of gliomas

In order to gain further insight into the composition of the immune infiltrate in the glioma subgroups, we dissected out the myeloid component in the tumor suspension. We combined the LGG and GBM IDH1mut (herein called IDHmut) for the subsequent comparisons with the other two groups of IDHwt GBMs. We observed that the percentage of neutrophils (Fig. 2A), myeloid derived suppressor cells (MDSCs) (Fig. 2B) and macrophages (Fig. 2C) was increased in GBMwt _hi compared to both, IDHmut and GBMwt_lo gliomas. We also analyzed the presence of CD206, a typical marker of alternatively activated (M2) macrophages. We found a higher proportion of CD11b+CD206+ cells in GBMwt_lo and GBMwt_hi, compared to IDHmut gliomas (Fig. 2D). Our panel was not designed to recognize specifically resident microglia, but we found no differences in the transcription of *P2RY12*, which is highly expressed on microglia, between the three subgroups (Supplementary Fig. 3A). Moreover, ionized calcium binding adaptor molecule 1 (IBA1) positive cells were detected in high proportion in all the tumors analyzed (Supplementary Fig. 3B). The number of microglial cells (Supplementary Fig. 3C), as well as the total amount of IBA1 protein (Supplementary Fig. 3D, E), was very homogenous among the different gliomas. These results suggest that the main differences in the immune compartment of the distinct glioma subgroups are due to the entrance of cells from the blood. Notably, the ratio of myeloid to lymphoid cells was lower in GBMwt_hi compared to the other two subgroups (Fig. 2E), suggesting that T cells infiltrate specially this subgroup of gliomas. In agreement with that, we observed that the proportion of T cells (CD3+) (Fig. 2F), in particular the CD4+ subset (Fig. 2G), was higher in GBMwt_hi tumors than in the other two subclasses. However, there was an increase in the percentage of CD3+ cells in GBMwt_lo compared to IDHmut gliomas (Fig. 2F). Furthermore, there was no difference between the percentages of CD8+ cells between the two subgroups of GBMs (Fig. 2H), which was higher in both compared to IDHmut tumors. As a result, the CD4/CD8 ratio was lower in the GBMwt_lo compared to GBMwt_hi tumors (Fig. 2I). This ratio has been linked to the appropriate lymphocyte function in other types of cancer^15^. Moreover, the proportion of PD1+ cells, which labels T cell exhaustion, was higher in GBMwt_hi compared to GBMwt_lo gliomas (Fig. 2J), whereas the amount of regulatory T cells (Tregs) was similar in the two groups (Fig. 2K). By contrast IDHmut gliomas presented fewer exhausted T cells (Fig. 2J) and Tregs (Fig. 2K). Taken together, these findings highlight the important dissimilarities in the immune profile of IDHmut vs IDHwt gliomas and suggest that the GBMwt_lo subgroup resembles IDHmut gliomas in their percentage of myeloid cells. Besides, we found differences in the lymphocyte content and function between the two subgroups of GBMs.

**Figure 2.**
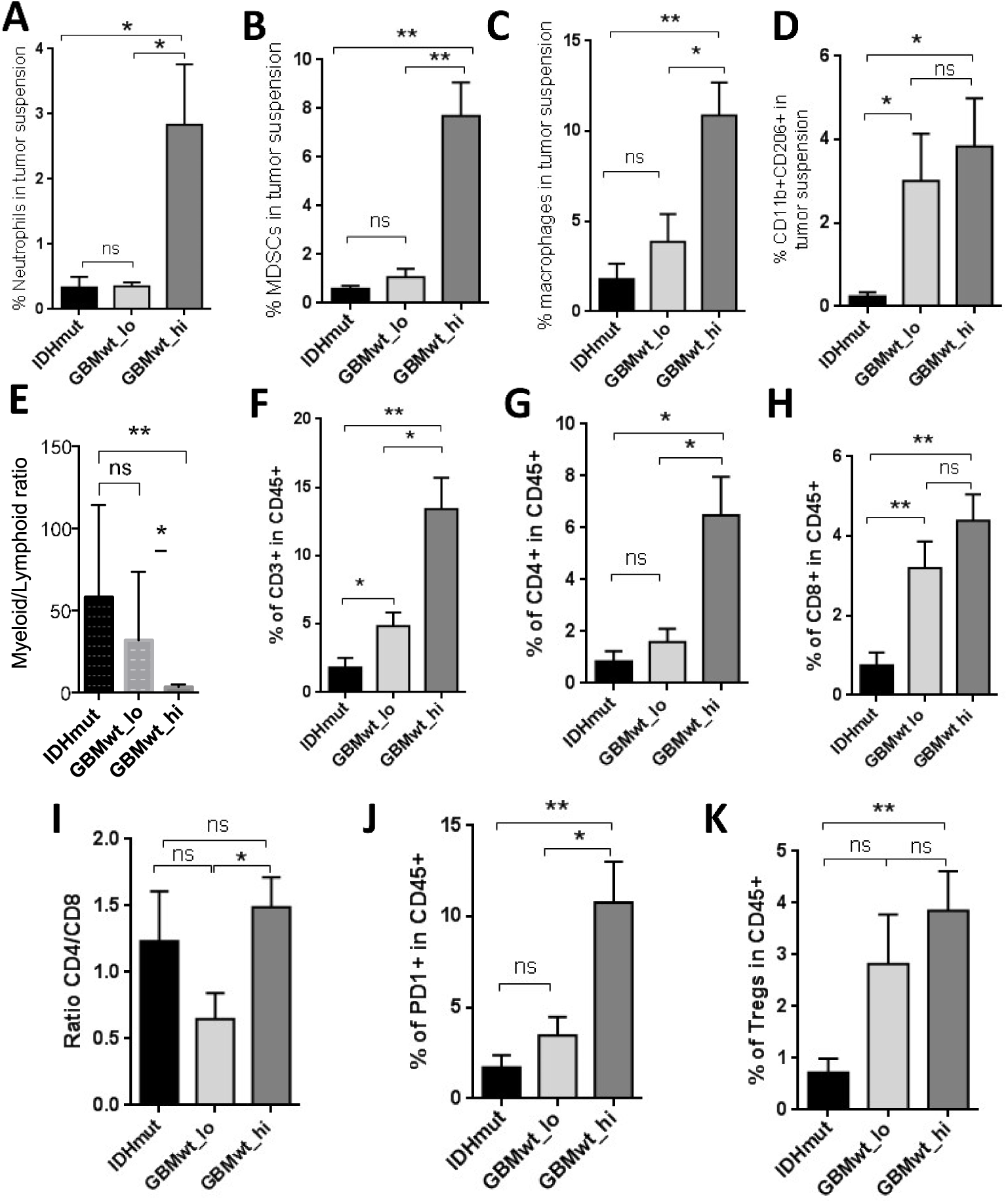
Proportions of myeloid and T cell subsets in glioma samples. (A-C) Percentage of neutrophils (A), MDSCs (B) and macrophages (C), on total tumor suspensions from IDHmut (IDHmut LGG and GBM) (n=6), GBMwt_lo (n=5), and GBMwt_hi (n=6) gliomas. (D-E) Percentage of CD206^+^ myeloid cells (D) and myeloid to lymphoid ratio (E) on total tumor suspensions from IDHmut (n=4), GBMwt_lo (n=6), and GBMwt_hi (n=10) gliomas. (F-I) Percentage of T cells (F), CD4^+^ T cells (G) and CD8^+^ T cells (H) and ratio of CD4^+^ to CD8^+^ T cells (I) within the CD45^+^ tumor subset of IDHmut (n=4), GBMwt_lo (n=6), and GBMwt_hi (n=10) gliomas. (J-K) Percentage of PD1^+^ exhausted T cells (J) and T regs (K) within the CD45^+^ tumor subset of IDHmut (n=5), GBMwt_lo (n=6), and GBMwt_hi (n=7) gliomas. *p<0.05, **p<0.01, ns: not significant.

### Enrichment of PD-L1 expression in the immune cells of highly infiltrated gliomas

Our flow cytometry analysis in gliomas revealed two levels of expression of PD-L1 (herein defined as PD-L1_lo and PD-L1_hi) (Fig. 3A), both in tumor (CD45-) and in immune (CD45+) cells. The expression profile was similar in GBMwt_lo and IDHmut tumors and very different from the GBMwt_hi gliomas (Fig. 3B), which showed a strong increase in the percentage of CD45/PDL1 double positive cells. The increment was significant in both the PD-L1_hi (Fig. 3C) and the PD-L1_lo (Fig. 3D) populations. Notably, there were no differences in the amount of tumor cells expressing high levels of PD-L1 among the different subgroups of gliomas (Fig. 3E). Altogether, our data suggest that there is a subgroup of IDHwt GBMs that contain a high immune infiltrate, enriched in myeloid cells and with a strong immunosuppressive profile: high content of MDSCs, CD206+ myeloid and PD-L1+ cells.

**Figure 3.**
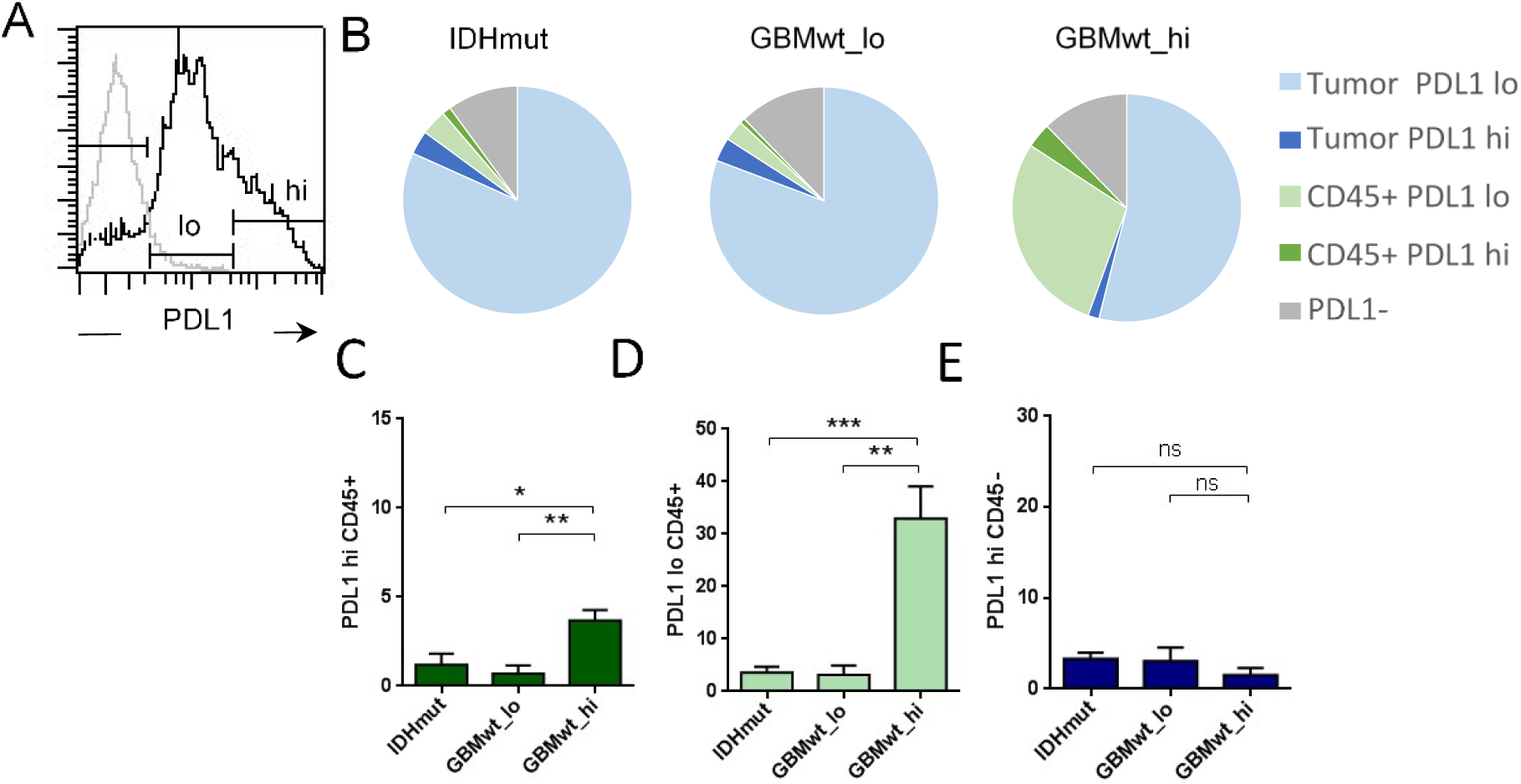
PD-L1 expression in gliomas. (A) The histogram shows the isotype control labelling in grey line (negative) and low (lo) and high (hi) levels of expression of PD-L1 in the total tumor suspension of a representative GBMwt_hi tumor. (B) The tart diagrams the percentage of PD-L1^lo^ and PD-L1^hi^ tumor cells (CD45^-^) and leukocytes (CD45^+^) on total tumor suspensions in each glioma subgroup: IDHmut (n=6), GBMwt_lo (n=4) GBMwt_hi (n=5). (C-E) Percentage of PD-L1^hi^CD45^+^ (C), PD-L1^lo^CD45^+^ (D) and PD-L1^hi^CD45^-^ (E) in each glioma subgroup. * p<0.05, **p<0.01, ***p<0.001

### The immune stratification of the tumors correlates with different vascular phenotypes

It has been proposed that the three different transcriptomic subtypes of gliomas (Proneural, PN, Classical, CL and Mesenchymal, MES) are associated with a different immune microenvironment^7^. When we analyzed our cohort of gliomas using qRT-PCR we noticed that, as expected, PN and MES transcripts were enriched (Supplementary Fig. 4A) and diminished (Supplementary Fig. 4B), respectively, in IDHmut gliomas compared to their wild-type counterparts. However, there were no differences in the expression of PN (Supplementary Fig. 4A) or MES (Supplementary Fig. 4B) markers between the two subgroups of IDHwt GBMs. Therefore, neither the genetic (Supplementary Fig. 2A) nor the expression profiles seem to explain the existence of two distinct immune patterns in the aggressive IDHwt gliomas.

In order to find disparities between the two subgroup of IDHwt GBMs that could correlate with their distinct immune landscapes we performed a macroscopic analysis of the tumors. Preoperative magnetic resonance imaging (MRI) revealed clear differences between IDHmut and IDHwt gliomas, especially in the contrast enhanced sequences (Fig. 4A)^16^. However, the T1+C and T2 images of GBMwt_lo tumors were very similar to the ones obtained in the immune-high GBM subgroup (Fig. 4A). By contrast, the macroscopic analysis of different vascular features revealed that the blood vasculature score (unbiased annotations from neurosurgeons) and the number of vessels with a large lumen (herein called dilated blood vessels (BVs)) were higher in the GBMwt_hi tumors (Fig. 4B and Supplementary Fig. 5). The IHC labelling of endothelial cells confirmed that the number of dilated vessels (Fig. 4C), as well as the CD34 density (Fig. 4D), correlated with the percentage of CD45+ cells measured by flow cytometry.

**Figure 4.**
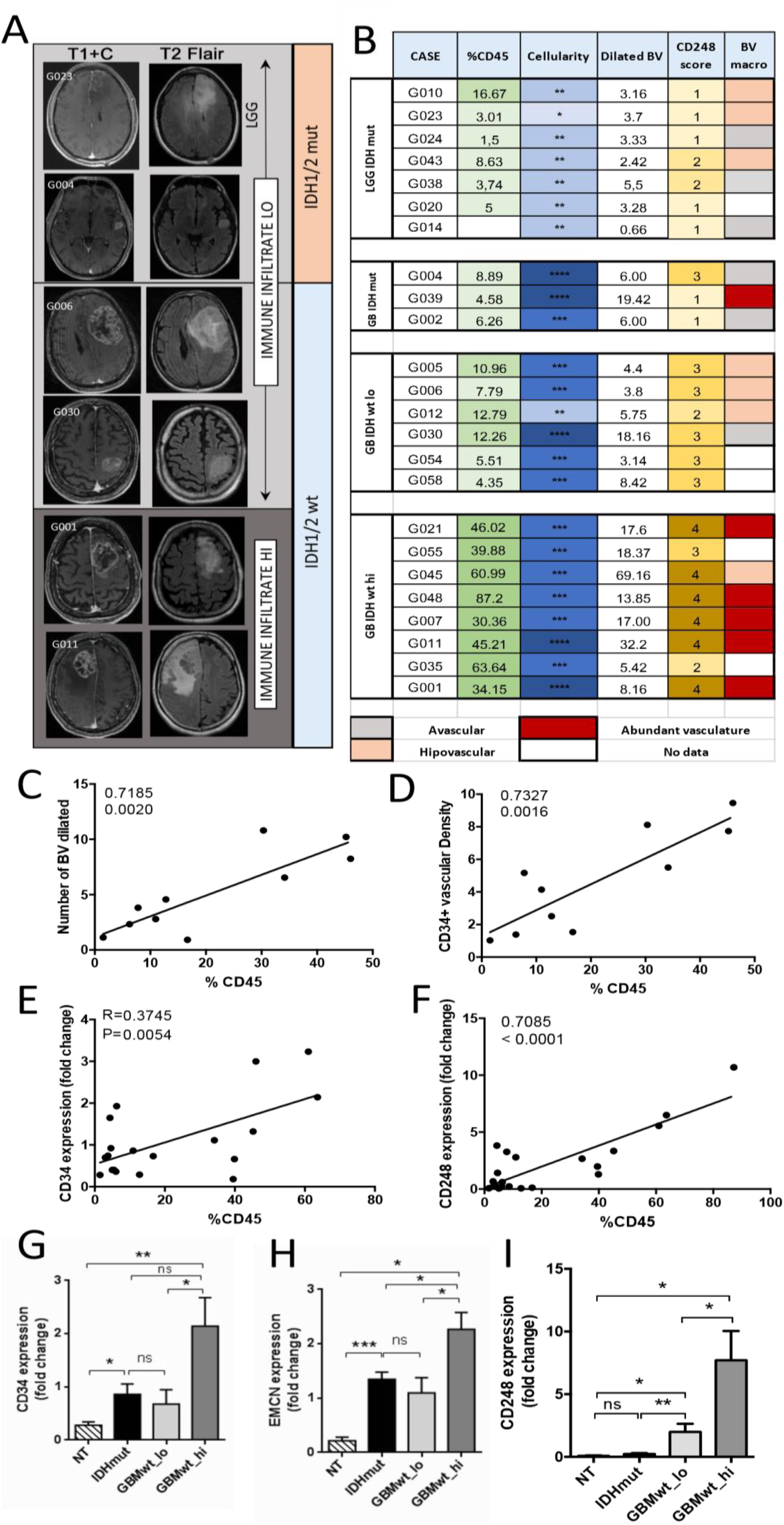
Correlation between the immune infiltrate and the vascular alterations in gliomas. (A) Representative MRIs (T1 post contrast (T1+C) and T2 FLAIR) of gliomas with low (LO) and high (HI) immune infiltrate. (B) The table aligns the percentage of CD45^+^ cells in each tumor with the cellularity (measured in the Hematoxilin & Eosine stainings), two vascular parameters measured by IHC (number of dilated blood vessels (BV) and CD248 score), and macroscopic evaluation of the tumor vascularization (BV macro) during surgery. (C-D) Correlation between the percentage of CD45^+^ cells and the number of dilated BVs (C) and the density of CD34 staining in the tissue sections (n=10). (E-F) Correlation between the percentage of CD45^+^ cells and *CD34* (E) and *CD248* (F) transcription measured by qRT-PCR analysis (n=21). (G-I) qRT-PCR analysis of *CD34* (G), *EMCN* (H), and *CD248* (I) expression in the different glioma subgroups: IDHmut (n=9), GBMwt_lo (n=6) and GBMwt_hi (n=11). *HPRT* transcription was used for normalization. * p<0.05, **p<0.01, ***p<0.001

To obtain an independent confirmation of these results, we performed a qRT-PCR analysis. We found a strong correlation between the CD45 content and the expression of *CD248* (Fig. 4E), which is highly expressed by tumor-pericytes in gliomas^17^, as well as with the expression of the endothelial *CD34* (Fig. 4F). Moreover, the transcription of *CD34* (Fig. 4G), *EMCN* (another marker of endothelial tumor cells) (Fig. 4H) and *CD248* (Fig. 4I) were increased in the GBMwt_hi group compared to the rest of gliomas. Notably, only the expression of *CD248*, was increased in GBMwt_lo tumors compared to IDHmut gliomas (Fig. 4I), which correlated with the higher CD248 score measured by IHC (Fig. 4B and Supplementary Fig. 5), suggesting a direct correlation between the absence of IDH mutations and the increase in tumor pericytes, as we have recently described^11^.

### Inverse correlation of the immune content with Tau expression

We have recently described that Tau, a protein associated with neurodegenerative diseases, is also expressed in glioma cells, particularly in the less aggressive tumors, where it obstructs glioma progression by blocking the formation of novel and aberrant tumor vessels. These effects were associated with a limited capacity of the gliomas cells to contribute to the pool of pericytes in Tau-high tumors, which results in a reduced amount of dilated BVs and a less efficient fueling of tumor growth^11^. We measured the amount of Tau in our cohort of gliomas and we observed that it accumulates in IDHmut gliomas (Fig. 5A-B). This result was not surprising given that the *MAPT/Tau* gene is epigenetically induced by the presence of mutant IDH proteins^11^. However, we also found an enrichment of Tau in GBMwt_lo compared to GBMwt_hi tumors (Fig. 5A-B), suggesting a possible correlation between this protein and the immune landscapes of gliomas. To test this hypothesis, we performed an in silico analysis of the TCGA database, which revealed a strong inverse correlation between the amount of *MAPT/Tau* transcription and overall survival or the expression of vascular-(*CD34* and *CD248*) (Fig. 5C and Supplementary Fig. 6A-B) and immune-(*CD3, CD4, CD11b* and *CD68*) Fig. 5C and Supplementary Fig. 6C-F) related genes. Notably, the transcript levels of *MAPT/Tau* and *CD248* were inversely and directly correlated, respectively, with several of the signatures associated with different immune cell populations (Fig. 5C) and with the inflammatory- and cytokine-related pathways (Fig. 5D). We also analyzed which genes were downregulated in Tau-high gliomas and we found that many of them were linked to the immune response (Supplementary Fig. 6G).

**Figure 5.**
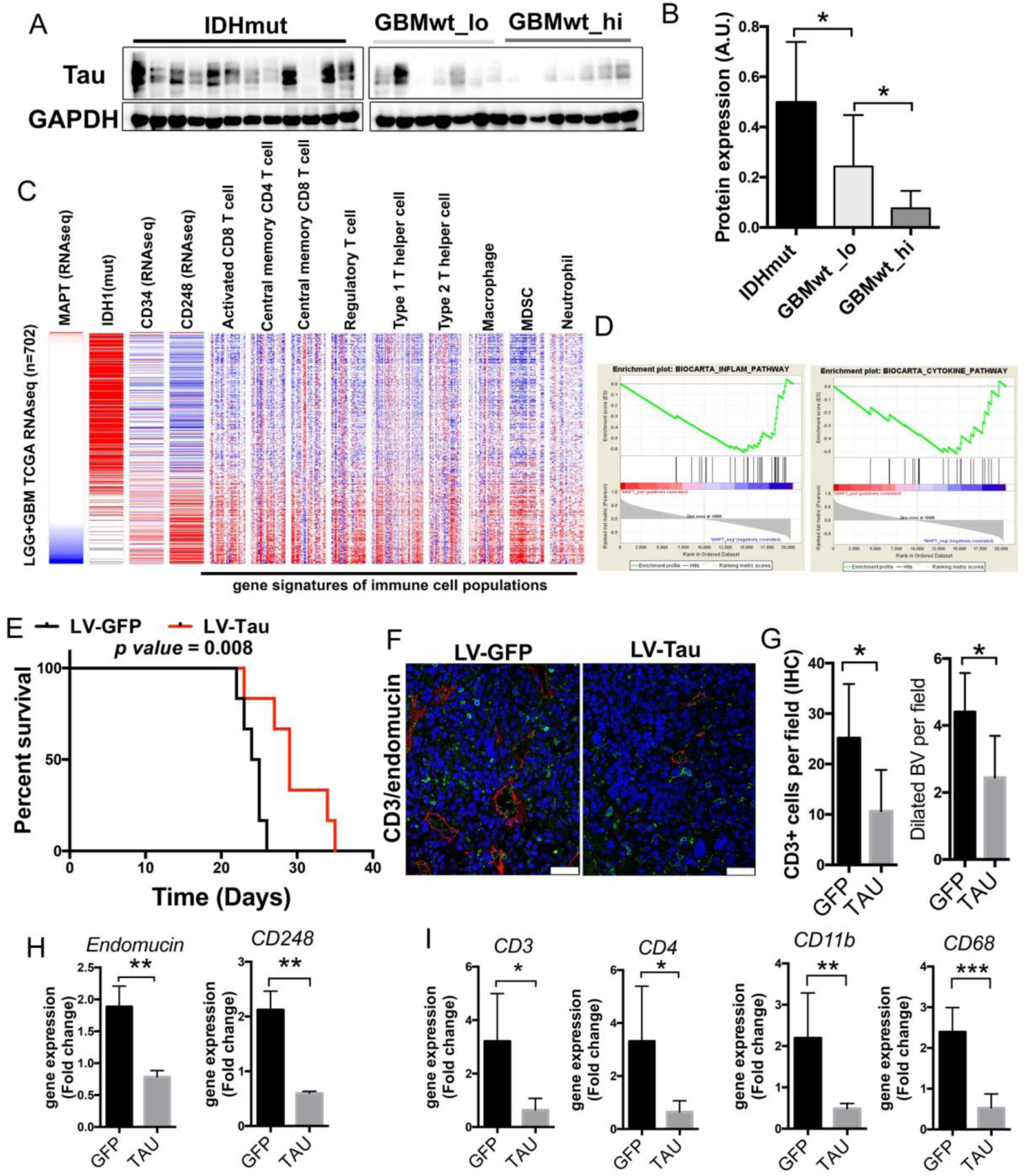
Tau expression correlates inversely with the immune content in gliomas. (A) WB analysis of Tau expression in extracts from IDHmut, GBMwt_lo and GBMwt_hi tumors. GAPDH level was used as a loading control. (B) Quantification of the WB in (A) (n=26). (C) Expression (RNAseq) of *IDH1, CD34, CD248* and 9 different immune populations signatures in gliomas (TCGA cohort). Samples were arranged according to their levels of expression of *MAPT/Tau* (n=702) (D) GSEA enrichment plot analysis using Tau gene expression values as template and the inflammatory and cytokine pathway gene-set from the Biocarta pathways database. (E) Kaplan-Meier overall survival curves of C57Bl/6 mice that were orthotopically injected with GL261 cells that overexpressed Tau or GFP proteins (n= 5). (F) Representative pictures of CD3 (green) and endomucin (red) staining in tumors from (E). (G) Quantification of the number of CD3^+^ cells and the number of dilated BVs per field in (F). (H-I) Expression of vascular (H) and immune (I) genes in the GL261 tumors (F). (n=4). Scale bar: 10μm. *p<0.05, **p<0.01, ***p<0.001

To get further insight we overexpressed Tau in GL261 cells, a well-known mouse glioma model. Tau overexpression increased the survival of mice bearing orthotopic tumors (Fig. 5E), which is in agreement with the increased survival of patients with low *MAPT/Tau* expression (Supplementary Fig. 6H). The analysis of the tumors revealed a decrease in the amount of infiltrating CD3 lymphocytes, in parallel with a reduction in the number of dilated BVs in the Tau-overexpressing gliomas (Fig. 5F-G). The transcriptomic analysis of the tumor tissues confirmed the inhibition of the expression of vascular- (Fig. 5H) and immune- (Fig. 5I) related genes in GL261-Tau tumors, compared to their control counterparts. These findings suggest that Tau modulates both, the immune phenotype and the vascular features of gliomas.

We have previously shown that the Tau expression is induced by IDH mutations and repressed by wild-type IDH1^11^, which is upregulated in primary GBs and promotes aggressive growth and therapy resistance^18^. Moreover, it has been shown that the expression of wild-type IDH1 reshapes the methylome and also affects gene expression^19^. In agreement with these data, we found that total *IDH1* expression was upregulated in those GBMs with a higher immune content (Supplementary Fig. 7A). This result suggests an explanation for the downregulation of Tau expression in the GBMwt_hi subgroup, which could be responsible, at least in part, for the increase in the vascular abnormalities and with the immune-enriched TME observed in these gliomas. However, we cannot discard that epigenetic changes induced by a higher amount of wild-type IDH could be affecting the expression of other immunomodulatory molecules as well. One such gene could be *HLA-A*, whose expression can be modulated by epigenetic mechanisms^20^. As a matter of fact, we observed a decrease in the amount of *HLA-A* transcription in the G-CIMP gliomas (Supplementary Fig. 7B), a phenotype associated with the presence of IDH mutations. When we analyzed our cohort, we observed that *HLA-A* transcription was elevated in the GBMwt_hi subgroup, in comparison with the rest of gliomas (Supplementary Fig. 7C). Moreover, we found a strong correlation between the expression of *HLA-A* and *IDH1*wt in the TCGA dataset (Supplementary Fig. 7D), similar to the one observed between *MAPT/*Tau and *IDH1*. Taken together, our results suggest that the balance between mutant and wild-type IDH function in gliomas is controlling the expression of Tau, and probably other proteins, to shape the vascular and the immune niche of gliomas.

## Discussion

A recent pan-cancer immunogenomic analysis have emphasized the unique microenvironment of glioma with an enrichment of the lymphocyte-depleted and the macrophage-enriched signatures in GBMs, whereas LGGs showed an immunologically quiet (“cold”) expression pattern^4^. Our detailed characterization of the immune content of different gliomas suggests that the presence of *IDH1/2* mutations, even more than the histological grading, is the best predictor of a reduced immune infiltration in gliomas. This observation is in agreement with the results obtained in mouse models^14^ and with the retrospective analysis of human data^7,8,21^. The paucity of immune cells in IDHmut gliomas could participate in the reduced aggressiveness of IDHmut gliomas, as leukocytes facilitates tumor proliferation^22^. As a drawback, tumors bearing IDH mutations could have an inferior response to immunotherapy. For that reason, several clinical trials have been designed combining inhibitors of mutant IDH with ICIs and vaccinations^23^.

GBM, even if we exclude IDHmut tumors, is not a homogenous entity. Here, we have described the existence of at least two different immune landscapes in IDHwt GBMs. Notably, the expression of PN or MES markers did not allow us to distinguish between these two immune subtypes of GBMs. These results differ from some computational^7^ and IHC^9,10^ studies, showing the enrichment of myeloid and lymphoid cells in the MES subtype. Another study, however, observed an accumulation of CL tumors among GBMs with a high immune infiltrate^8^.

Regarding clinical implications, there are evidences of a possible correlation between the presence of innate immunity cells and the aggressive behavior of gliomas^6^. However, our survival analysis could not detect significant differences in the clinical behavior of the two IDHwt GBM subgroups, as other investigators have shown^21^. However, our study is conditioned by the limited number of cases to test survival differences.

The immune compartment of the GBMwt_high subgroup can account for half of the tumor mass, basically at the expense of recruited cells. Notably, no clear differences were found in the amount of microglia amid the different subgroups. Our results are in agreement with recently published data showing that the main immune signature in IDHmut gliomas correspond to microglia, whereas in the wild-type tumors there is an accumulation of infiltrating myeloid and T cells^24^. Notably, we have found a high level of expression of CD206, a marker of pro-tumoral M2 macrophages, in myeloid cells in the GBMwt_high tumors, and a strong increase in PD-L1 expression between this subgroup and the rest of gliomas. This increment is mostly due to its presence on the surface of the immune cells, where myeloid cells are enriched. Although we cannot discard the presence of PD-L1 in lymphocytes^25^, its expression has already been described in GBM infiltrating macrophages^26^. Based on our results, strategies to impair the high immunosuppressive environment of GBMwt_hi tumors are essential. In agreement with that, it has been recently reported that targeting of myeloid cells increases the response to anti-PD-1 in glioma mouse models^27^. By contrast, the low PD-L1 expression in the rest of gliomas could be hampering the result of antibodies targeting this molecule. However, the lower CD4/CD8 ratio and the scarcity of myeloid cells in the GBMwt_lo gliomas could make them more prone to respond to different immunotherapies. In any case, our results highlight the relevance of a patient selection, reasonably based on the vascular-immune profile, to improve the success of future immunotherapies.

Several groups have attributed the scarcity of T cells in IDHmut gliomas to the accumulation of the oncometabolite 2-hydroxyglutarate^14,23^. However, these mutations could be affecting other components of the TME, specially macrophages and MDSCs through different mechanisms, such as implementing an specific cytokine program as it has been discovered in IDH mutant gliomas^28^. Here, we propose that, secondary to IDH mutations and/or to a reduction in the amount of wild-type IDH enzymes, Tau accumulates in the tumors. By contrast, GBMwt_hi gliomas, which contain the higher expression of IDHwt, had a reduced amount of Tau. It is important to point out that the balance of wild-type and mutant IDH proteins controls the clinical outcome of gliomas, including their sensitivity to radiation and chemotherapy^29^. In agreement with our previous results^11^, we have shown here that as the level of Tau decreases, the number of pericytes increases and the tumor vasculature appears distorted, with numerous enlarged vessels. Moreover, we have observed a striking positive correlation between the presence of the immune infiltrate and the appearance of these vascular abnormalities. Additionally, and in agreement with our hypothesis, the levels of Tau decreased in parallel to these changes in the immune profiles. Furthermore, the analysis of different orthotopic mouse and human glioma models have confirmed that overexpression of Tau reduces the amount of both leukocytes and myeloid cells in the tumors, in parallel to changes in the cytokine and inflammatory signatures as well as with the normalization of the vessels. Our results underline the intricate connection between the two main components of the glioma niche. In relation with that, the main angiogenic factor, vascular endothelial growth factor (VEGF), can inhibit the function of T cells and increase the recruitment of regulatory Tregs and MDSCs^30^. Moreover, it has been shown that pericytes can support tumor growth via immunsuppression^31^. However, once in the tumors, the immune cells can also induce changes in the vascular compartment as they have a strong pro-angiogenic role^32,33^. Our data does not allow us to discriminate which came first: the chicken or the egg. In the presence of low levels of Tau, an increase in the number of glioma-derived-pericytes and the subsequent changes in the BBB, could directly ease the extravasation of hematopoietic cells. However, we cannot discard that the downregulation of Tau might inhibit directly the secretion of molecules that attract myeloid cells and this could further contribute to the vascular phenotype. Moreover, other proteins epigenetically induced or repressed by the balance between mutant and wild-type IDH will probably participate in the formation of the different immune-vascular landscapes. In any case, our results suggest that combined strategies targeting these two components of the tumors’stroma might lead to promising results, as it is already being tested in recurrent GBMs, using nivolumab and bevacizumab (NCT03452579, NCT03743662, NCT03890952). However, based on our results, an effort should be made to understand which group of patients could benefit most from these combinatorial treatments and also to design new treatment schemes.

## Acknowledgments

The authors would like to acknowledge the Confocal and the Animal Facility service personnel, for their technical support.

## References

1. Louis DN, Perry A, Reifenberger G, et al. The 2016 World Health Organization Classification of Tumors of the Central Nervous System: a summary. Acta Neuropathol. 2016; 131(6):803–820.

2. Filley AC, Henriquez M, Dey M. Recurrent glioma clinical trial, CheckMate-143: the game is not over yet. Oncotarget. 2017; 8(53):91779–91794.

3. Cloughesy TF, Mochizuki AY, Orpilla JR, et al. Neoadjuvant anti-PD-1 immunotherapy promotes a survival benefit with intratumoral and systemic immune responses in recurrent glioblastoma. Nat.Med. 2019; 25(3):477–486.

4. Thorsson V, Gibbs DL, Brown SD, et al. The Immune Landscape of Cancer. Immunity. 2018; 48(4):812–830.

5. Pombo Antunes AR, Scheyltjens I, Duerinck J, Neyns B, Movahedi K, Van Ginderachter JA. Understanding the glioblastoma immune microenvironment as basis for the development of new immunotherapeutic strategies. Elife. 2020; 9.

6. Gieryng A, Pszczolkowska D, Walentynowicz KA, Rajan WD, Kaminska B. Immune microenvironment of gliomas. Lab Invest. 2017; 97(5):498–518.

7. Wang Q, Hu B, Hu X, et al. Tumor Evolution of Glioma-Intrinsic Gene Expression Subtypes Associates with Immunological Changes in the Microenvironment. Cancer Cell. 2017; 32(1):42–56.

8. Luoto S, Hermelo I, Vuorinen EM, et al. Computational Characterization of Suppressive Immune Microenvironments in Glioblastoma. Cancer Res. 2018; 78(19):5574–5585.

9. Martinez-Lage M, Lynch TM, Bi Y, et al. Immune landscapes associated with different glioblastoma molecular subtypes. Acta Neuropathol.Commun. 2019; 7(1):203.

10. Kaffes I, Szulzewsky F, Chen Z, et al. Human Mesenchymal glioblastomas are characterized by an increased immune cell presence compared to Proneural and Classical tumors. Oncoimmunology. 2019; 8(11):e1655360.

11. Gargini R, Segura-Collar B, Herranz B, et al. The IDH-TAU-EGFR triad defines the neovascular landscape of diffuse gliomas. Sci.Transl.Med. 2020; 12(527).

12. Cantero D, Rodriguez de LA, Moreno dlP, et al. Molecular Study of Long-Term Survivors of Glioblastoma by Gene-Targeted NGS. J.Neuropathol.Exp.Neurol. 2018; 77(8):710–716.

13. Charoentong P, Finotello F, Angelova M, et al. Pan-cancer Immunogenomic Analyses Reveal Genotype-Immunophenotype Relationships and Predictors of Response to Checkpoint Blockade. Cell Rep. 2017; 18(1):248–262.

14. Amankulor NM, Kim Y, Arora S, et al. Mutant IDH1 regulates the tumor-associated immune system in gliomas. Genes Dev. 2017; 31(8):774–786.

15. Shah W, Yan X, Jing L, Zhou Y, Chen H, Wang Y. A reversed CD4/CD8 ratio of tumor-infiltrating lymphocytes and a high percentage of CD4(+)FOXP3(+) regulatory T cells are significantly associated with clinical outcome in squamous cell carcinoma of the cervix. Cell Mol Immunol. 2011; 8(1):59–66.

16. Lai A, Kharbanda S, Pope WB, et al. Evidence for sequenced molecular evolution of IDH1 mutant glioblastoma from a distinct cell of origin. J.Clin.Oncol. 2011; 29(34):4482–4490.

17. Simonavicius N, Robertson D, Bax DA, Jones C, Huijbers IJ, Isacke CM. Endosialin (CD248) is a marker of tumor-associated pericytes in high-grade glioma. Mod.Pathol. 2008; 21(3):308–315.

18. Calvert AE, Chalastanis A, Wu Y, et al. Cancer-Associated IDH1 Promotes Growth and Resistance to Targeted Therapies in the Absence of Mutation. Cell Rep. 2017; 19(9):1858–1873.

19. Turcan S, Rohle D, Goenka A, et al. IDH1 mutation is sufficient to establish the glioma hypermethylator phenotype. Nature. 2012; 483(7390):479–483.

20. Berghoff AS, Kiesel B, Widhalm G, et al. Programmed death ligand 1 expression and tumor-infiltrating lymphocytes in glioblastoma. Neuro Oncol. 2015; 17(8):1064–1075.

21. Jeanmougin M, Håvik AB, Cekaite L, et al. Improved prognostication of glioblastoma beyond molecular subtyping by transcriptional profiling of the tumor microenvironment. Mol Oncol. 2020; 14(5):1016–1027.

22. Hambardzumyan D, Gutmann DH, Kettenmann H. The role of microglia and macrophages in glioma maintenance and progression. Nat.Neurosci. 2016; 19(1):20–27.

23. Friedrich M, Bunse L, Wick W, Platten M. Perspectives of immunotherapy in isocitrate dehydrogenase-mutant gliomas. Curr Opin Oncol. 2018; 30(6):368–374.

24. Klemm F, Maas RR, Bowman RL, et al. Interrogation of the Microenvironmental Landscape in Brain Tumors Reveals Disease-Specific Alterations of Immune Cells. Cell. 2020; 181(7):1643–1660.e1617.

25. Nduom EK, Wei J, Yaghi NK, et al. PD-L1 expression and prognostic impact in glioblastoma. Neuro Oncol. 2016; 18(2):195–205.

26. Antonios JP, Soto H, Everson RG, et al. Immunosuppressive tumor-infiltrating myeloid cells mediate adaptive immune resistance via a PD-1/PD-L1 mechanism in glioblastoma. Neuro.Oncol. 2017; 19(6):796–807.

27. Flores-Toro JA, Luo D, Gopinath A, et al. CCR2 inhibition reduces tumor myeloid cells and unmasks a checkpoint inhibitor effect to slow progression of resistant murine gliomas. Proc.Natl.Acad.Sci.U.S.A. 2020; 117(2):1129–1138.

28. Venteicher AS, Tirosh I, Hebert C, et al. Decoupling genetics, lineages, and microenvironment in IDH-mutant gliomas by single-cell RNA-seq. Science. 2017; 355:1391.

29. Ichimura K, Narita Y, Hawkins CE. Diffusely infiltrating astrocytomas: pathology, molecular mechanisms and markers. Acta Neuropathol. 2015; 129(6):789–808.

30. Yang J, Yan J, Liu B. Targeting VEGF/VEGFR to Modulate Antitumor Immunity. Front Immunol. 2018; 9:978.

31. Sena IFG, Paiva AE, Prazeres P, et al. Glioblastoma-activated pericytes support tumor growth via immunosuppression. Cancer Med. 2018; 7(4):1232–1239.

32. Tian L, Goldstein A, Wang H, et al. Mutual regulation of tumour vessel normalization and immunostimulatory reprogramming. Nature. 2017; 544(7649):250–254.

33. Sidibe A, Ropraz P, Jemelin S, et al. Angiogenic factor-driven inflammation promotes extravasation of human proangiogenic monocytes to tumours. Nat.Commun. 2018; 9(1):355.

