## Supplemental Figures for "Immune profiling of gliomas reveals a connection with Tau function and the tumor vasculature"

**Supplementary Material:**

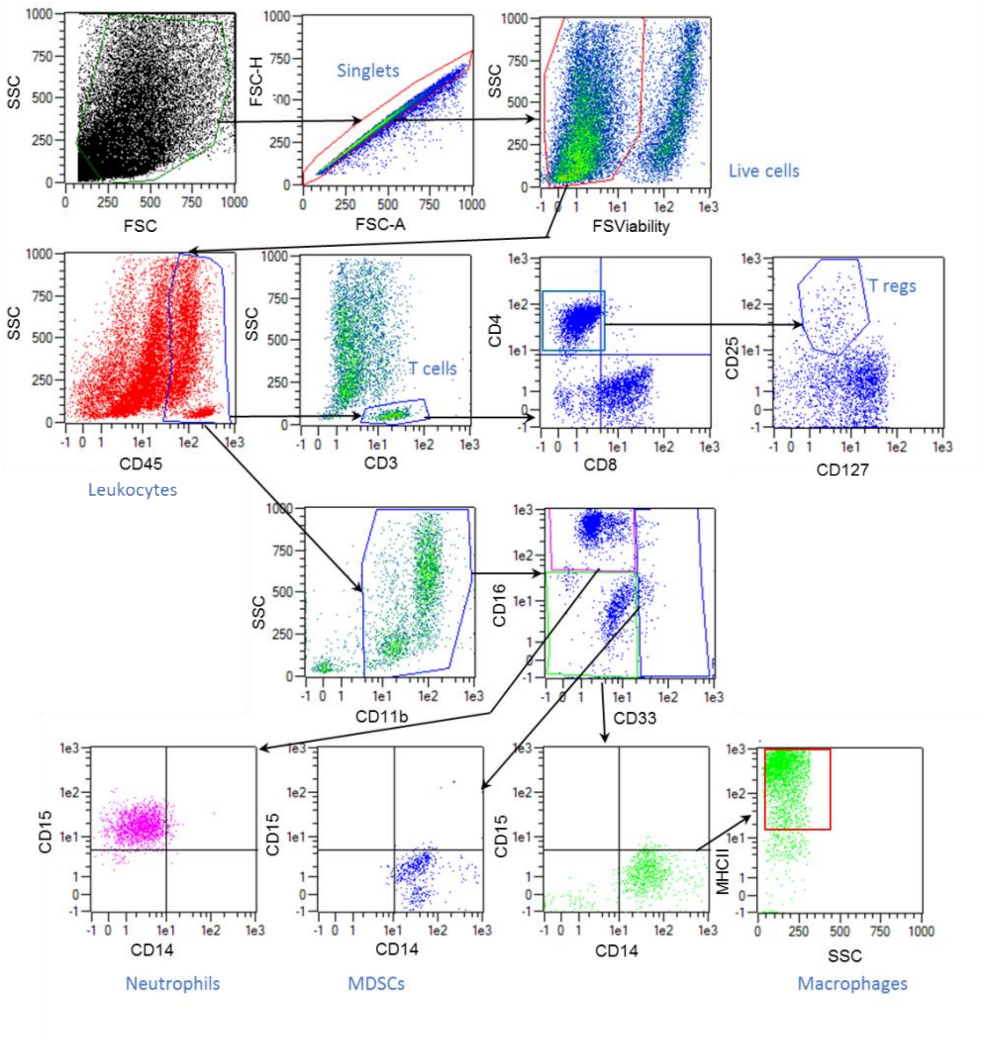


**Supplementary Figure 1.** Representative gating FACS plots to identify leukocytes and myeloid cells in human glioma tissues.

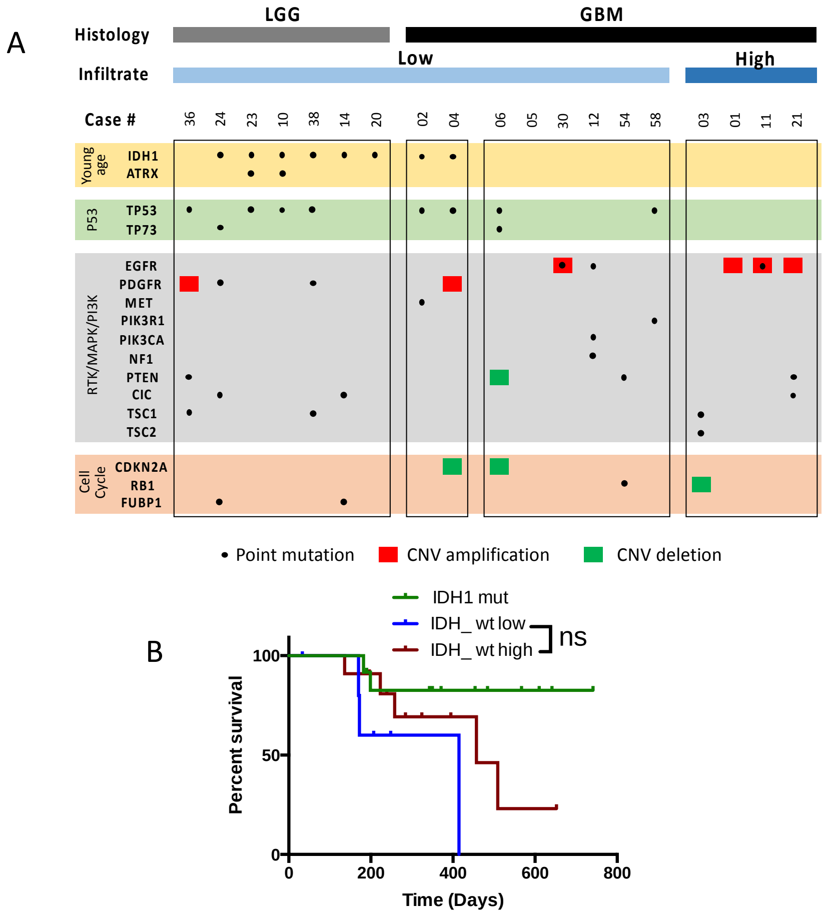


**Supplementary Figure 2.** Sequencing and Overall survival in de glioma cohort (A) Distribution of point mutations and copy number variations in 20 genes frequently altered in gliomas. (B) Kaplan-Meier survival curves of the three different groups associated with the immune phenotype, IDH mut, GBMwt_lo, GBMwt_hi gliomas.

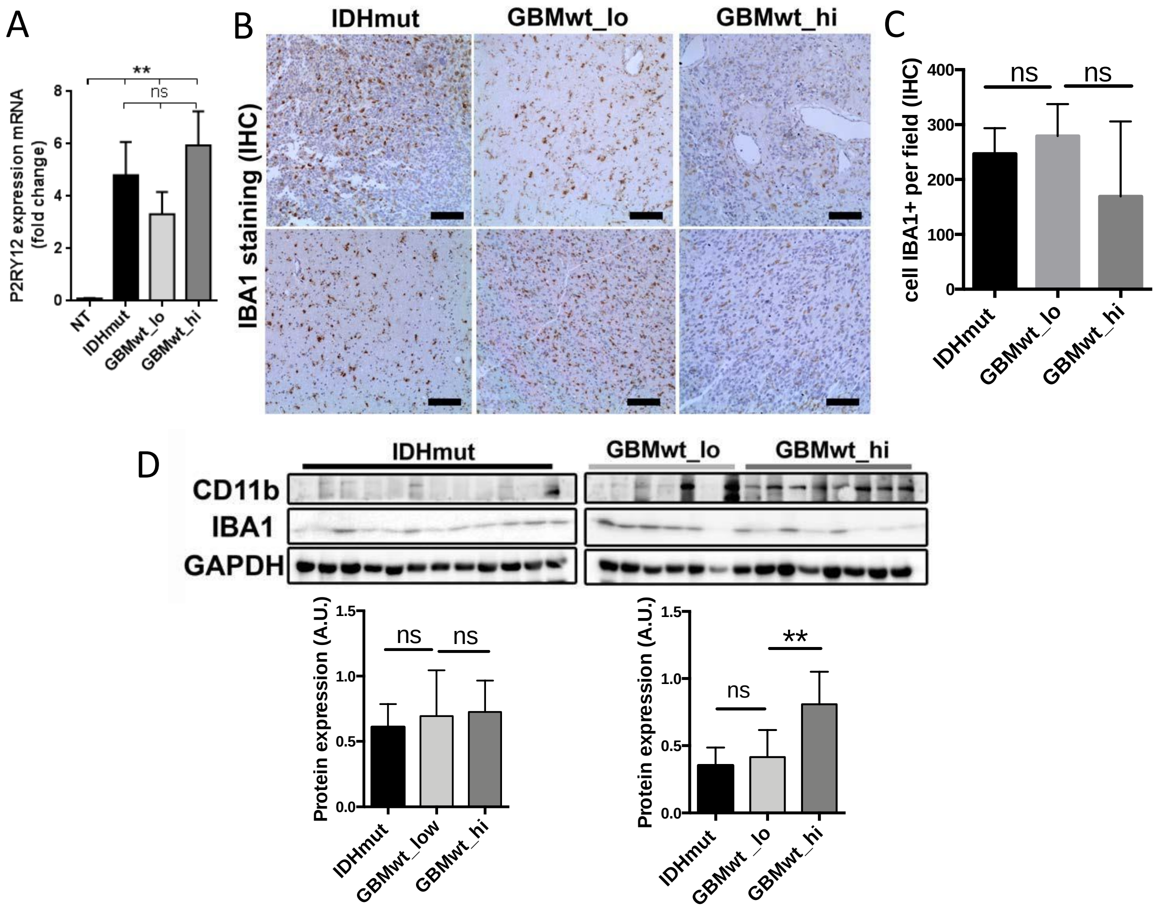


**Supplementary Figure 3.** Microglia content is similar in IDH mut, GBMwt_lo, GBMwt_hi gliomas. (A) Fold change values of qRT-PCR analysis of P2RY12 in tumor tissue from IDHmut, GBMwt_lo, GBMwt_hi (n=6). (B) Representative pictures obtained from IHQ for IBA1 in the three groups of gliomas. (C) Quantification of IBA1 levels from IHQ showed in B (n=6). (D) WB analysis of CD11b and IBA1 expression in tumor tissue extracts from IDHmut (black), GBMwt_lo (dark grey), GBMwt_hi (light grey) tumors. GAPDH level as a loading control. Quantification of levels of CD11b and IBA1 expression from WB showed in D. (**p<0.01, ns not significant)

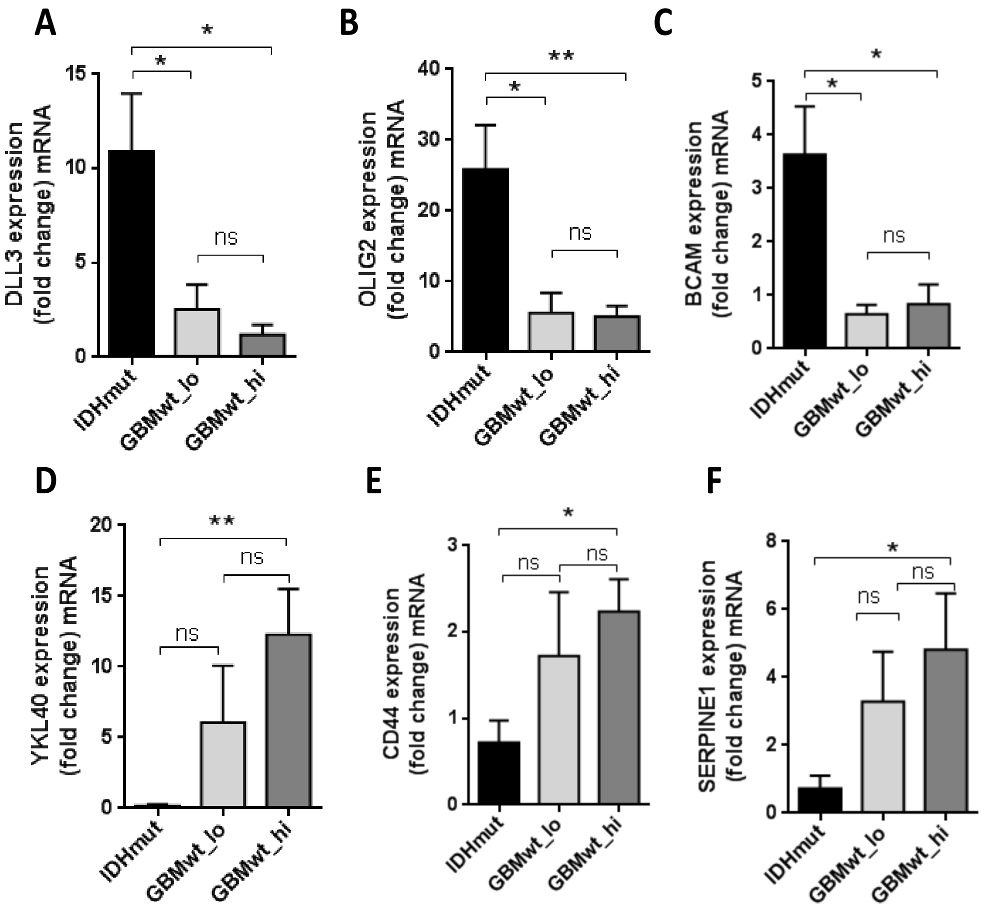


**Supplementary Figure 4.** Proneural or mesenquimal markers expression in IDHmut, GBMwt_lo, GBMwt_hi gliomas. Fold change values of qRT-PCR analysis of expression of (A-C) typical proneural markers as DLL3, OLIG2, BCAM. (D-F) typical mesenchymal markers as YKL40, CD44 or SERPINE1. (n=26). (* $p<0.05$, ** $p<0.01$, ns not significant)

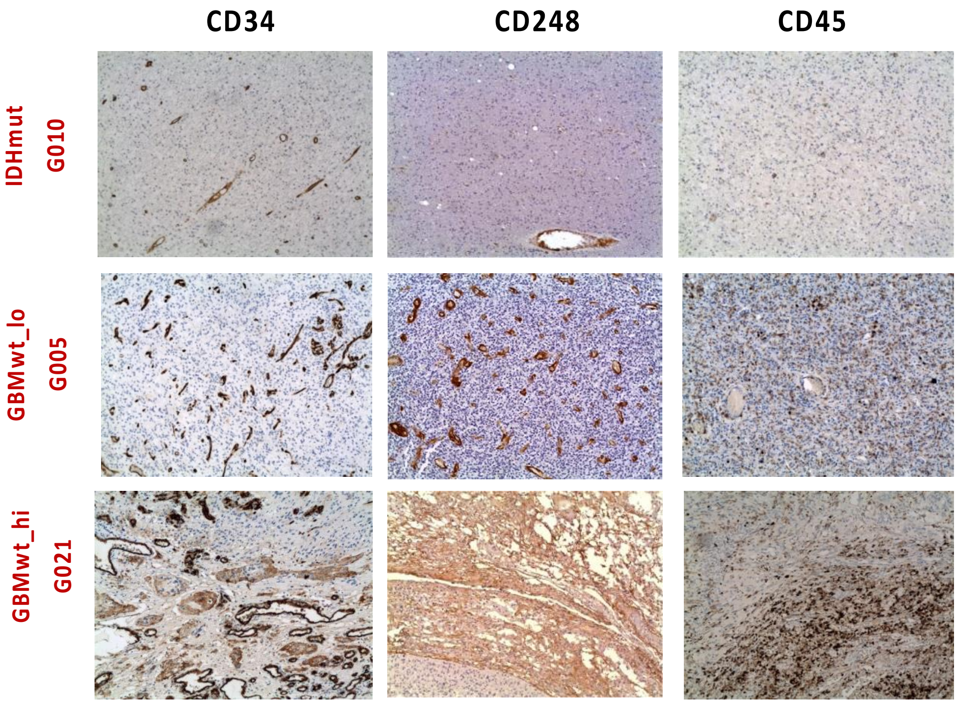


**Supplementary Figure 5.** Positive correlation between $CD45^{+}$ and $CD34^{+}$ or $CD248^{+}$ cells content in gliomas. Representative pictures from IHQs of CD45 (leukocyte marker), CD34 (endothelial marker) or CD248 (pericyte marker) in each group of gliomas.

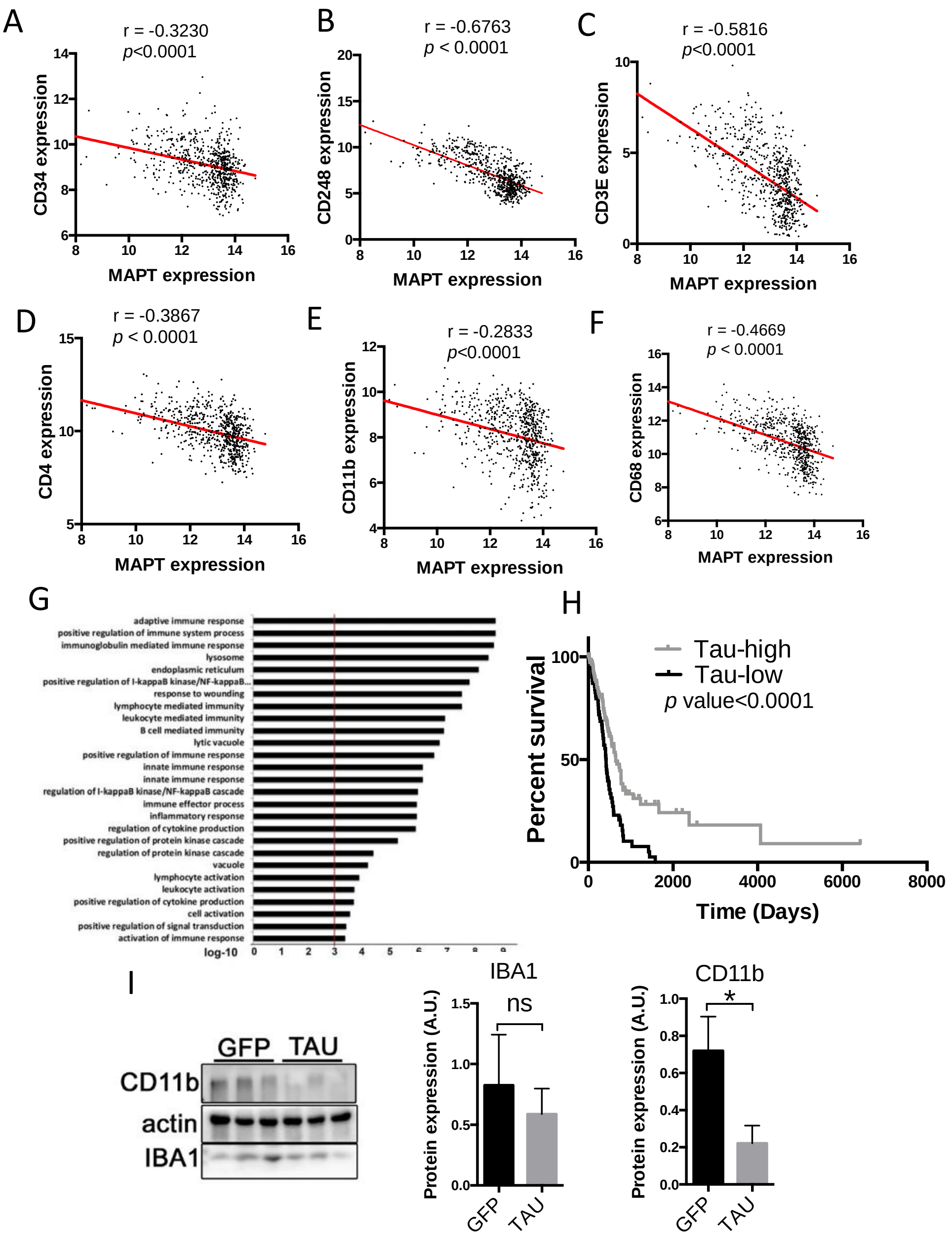


**Supplementary Figure 6.** Inverse correlation between Tau expression and vascular alterations or immune cells content in gliomas. Correlation of the mRNA expression by RNAseq of *Tau (MAPT)* with that of (A) CD34 (B) CD248 (C) CD3E (D) CD4 (E) CD11b and (F) CD68 in gliomas using the TCGA cohort (n=661). (G) Top enriched Gene Ontology (GO) biological process for the cluster of 500 genes that are negatively correlated with Tau (MAPT) in gliomas. We used the LGG+GBM merge cohort and the DAVID gene ontology program. (H) Kaplan-Meier overall survival curves of patients from the TCGA cohorte glioma IDH wt (n=191) and patients in each cohort were stratified into 2 groups based on high and low *Tau (MAPT)* expression values, log-rank (Mantel-Cox) test. (D) WB analysis of CD11b and IBA1 expression in tumor tissue extracts from GFP and Tau-GL261. GAPDH level as a loading control and quantification of levels of CD11b and IBA1 expression. (* p<0.05, **p<0.01, ns not significant).

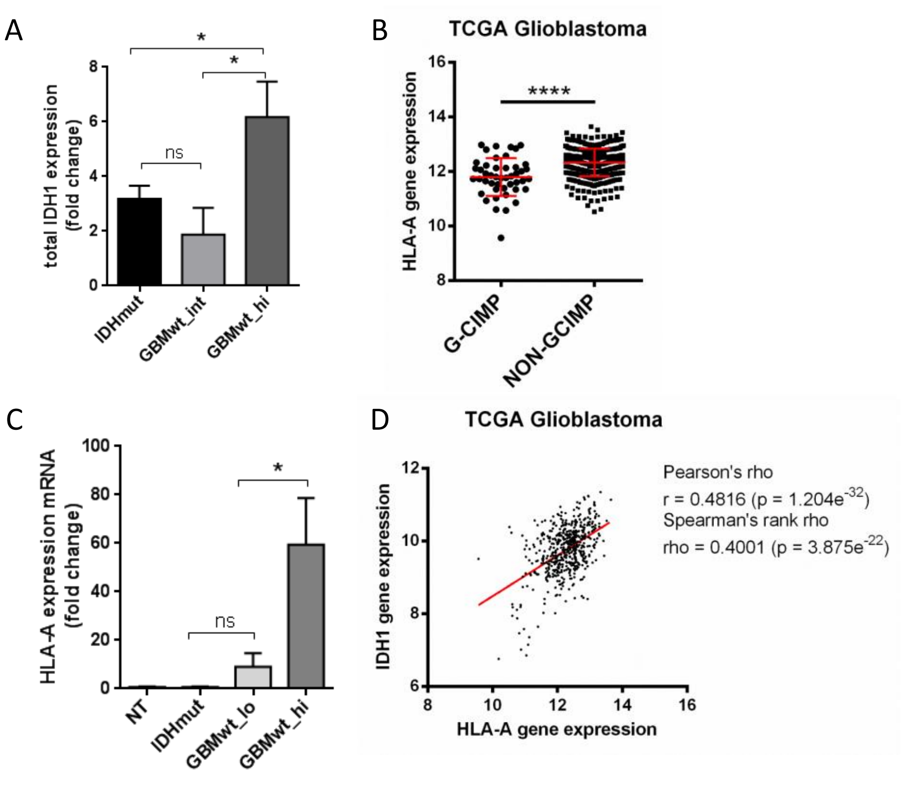


**Supplementary Figure 7.** Epigenetic changes could explain differences between the groups. **(**A) Fold change values of qRT-PCR analysis of IDH1 expression (n=6). (B) Analysis of HLA-A mRNA expression by RNAseq in gliomas (TCGA cohort) grouped according to their G-CIMP status (G-CIMP n=46 and NON-GCIMP n=475). (C) Fold change values of qRT-PCR analysis of HLA-A expression (n=6). (D) Correlation of the expression mRNA of IDH1 and HLA-A using TCGA glioma cohort (n=539). Paired T-test was performed (*p<0.05, ****p<0.0001, ns not significant).
