## Supplemental Table 1 for "Immune profiling of gliomas reveals a connection with Tau function and the tumor vasculature"

**Table 1. Study patients characteristics.**

| Patient | Age | Gender | Grade of resection | Diagnosis | Tumor Grade | *IDH* | *ATRX* | Immune Profile | *MGMT* | *TERT* | 1st line treatment | Time to PD | Time of F/U | Status after F/U |
|---|---|---|---|---|---|---|---|---|---|---|---|---|---|---|
| G001 | 82 | female | complete | astrocitoma | IV | WT | WT | GBMwt_hi | methylated | C228T | Stupp | 4.5 | 7.6 | death |
| G002 | 36 | female | complete | astrocitoma | IV | Mut | Mut | IDHmut or LGG | methylated | wt | Stupp | No PD | 25 | alive |
| G003 | 68 | male | complete | astrocitoma | IV | WT | Mut | GBMwt_hi | methylated | wt | Stupp | 8.2 | 8.9 | death |
| G004 | 42 | male | complete | astrocitoma | IV | Mut | Mut. | IDHmut or LGG | methylated | wt | Stupp | 4.3 | 5.5 | death |
| G005 | 70 | male | complete | astrocitoma | IV | WT | WT | GBMwt_lo | methylated | C228T | none | 4.6 | 4.6 | death |
| G006 | 57 | male | complete | astrocitoma | IV | WT | WT | GBMwt_lo | methylated | C228T | none | 1 | 1 | death |
| G007 | 34 | female | complete | astrocitoma | IV | WT | WT | GBMwt_hi | methylated | wt | Stupp | 16.7 | 23 | death |
| G010 | 30 | female | complete | astrocitoma | III | Mut | Mut | IDHmut or LGG | methylated | wt | Stupp | No PD | 22.4 | alive |
| G011 | 52 | male | complete | astrocitoma | IV | WT | WT | GBMwt_hi | unmethylated | C228T | Stupp | 17.3 | 21.6 | alive |
| G012 | 53 | male | partial | astrocitoma | IV | WT | WT | GBMwt_lo | methylated | C228T | Stupp | 6.6 | 13.6 | death |
| G014 | 30 | female | partial | oligodendroglioma | II | Mut | WT | IDHmut or LGG | methylated | C228T | RT+PCV | No PD | 20 | alive |
| G020 | 46 | male | complete | oligodendroglioma | II | Mut | WT | IDHmut or LGG | methylated | C228T | RT+PCV | No PD | 18.9 | alive |
| G021 | 55 | female | complete | astrocitoma | IV | WT | WT | GBMwt_hi | methylated | wt | Stupp | 6.6 | 14.7 | death |
| G023 | 45 | female | complete | astrocitoma | III | Mut | Mut | IDHmut or LGG | methylated | wt | Stupp | No PD | 14.9 | alive |
| G024 | 35 | female | complete | oligodendroglioma | II | Mut | WT | IDHmut or LGG | methylated | C228T | none | No PD | 16.8 | alive |
| G030 | 63 | male | complete | astrocitoma | IV | WT | WT | GBMwt_lo | methylated | C228T | Stupp | 5.5 | 5.5 | death |
| G035 | 65 | female | complete | astrocitoma | IV | WT | WT | GBMwt_hi | no data | C228T | Stupp | 4 | 14.2 | alive |
| G036 | 76 | female | partial | astrocitoma | III | WT | WT | IDHmut or LGG | methylated | C228T | temozolomide | 4.1 | 4.2 | death |
| G038 | 39 | male | complete | astrocitoma | III | Mut | Mut | IDHmut or LGG | no data | wt | Stupp | No PD | 13.3 | alive |
| G039 | 47 | male | complete | astrocitoma | IV | Mut | Mut. | IDHmut or LGG | methylated | wt | Stupp | No PD | 13 | alive |
| G043 | 38 | male | partial | astrocitoma | III | Mut | Mut. | IDHmut or LGG | methylated | C228T | Stupp | No PD | 12 | alive |
| G045 | 65 | male | complete | astrocitoma | IV | WT | WT | GBMwt_hi | no data | no data | Stupp | No PD | 11 | alive |
| G048 | 69 | female | complete | astrocitoma | IV | WT | WT | GBMwt_hi | no data | C228T | Stupp | 9.3 | 10.6 | alive |
| G054 | 42 | male | complete | astrocitoma | IV | WT | WT | GBMwt_lo | no data | wt | Stupp | No PD | 9.7 | alive |
| G055 | 71 | female | partial | astrocitoma | IV | WT | WT | GBMwt_hi | methylated | C228T | none | 4.4 | 4.4 | death |
| G058 | 70 | male | complete | astrocitoma | IV | WT | WT | GBMwt_lo | methylated | C250T | Stupp | No PD | 7.6 | alive |
| G059 | 50 | male | complete | astrocitoma | III | WT | WT | IDHmut or LGG | unmethylated | C228T | Stupp | No PD | 7.2 | alive |
| G060 | 50 | male | complete | astrocitoma | IV | WT | WT | GBMwt_hi | methylated | C228T | Stupp | No PD | 7.1 | alive |
