## Supplemental Table 2 for "Immune profiling of gliomas reveals a connection with Tau function and the tumor vasculature"

| Purpose | Antigen | Conjugated | Catalog number | Supplier | Clone |
|---|---|---|---|---|---|
| FACS | cd25 | 421 | 302629 | Biolegend | BC96 |
| | cd3 | vg | 563109 | BD | UCHT1 |
| | cd45 | 488 | 130-080-202 | Miltenyi | REA747 |
| | cd127 | pe | 561028 | BD | HIL-7R-M21 |
| | cd8 | pecy5 | 565310 | BD | SK1 |
| | cd4 | 647 | 300520 | Biolegend | RPA-T4 |
| | pd1 | 421 | 562516 | BD | EH12.1 |
| | cd16 | 421 | 302037 | Biolegend | 3G8 |
| | cd14 | pe | 130-110-577 | Miltenyi | REA599 |
| | cd15 | vg | 301910 | Biolegend | HI98 |
| | cd33 | pecy5 | 366615 | Biolegend | P67.6 |
| | cd11b | 647 | 130-098-087 | Miltenyi | M1/70 |
| | pdl1 | vg | 329713 | Biolegend | 29E.2A3 |
| | cd206 | pecy5 | 321121 | Biolegend | 15-2 |
| | mhcii | 421 | 562805 | BD | G46-6 |
| WB | CD11b | Purified | SAB1305652 | Sigma | 40TST |
| | IBA1 | Purified | ab16588 | Abcam | EPR 16588 |
| | GAPDH | Purified | sc-47724 | Santa cruz | 0411 |
| | Tau | Purified | 577801 | CALBIOCHEM | Tau-5 |
| IHC | CD34 | Purified | NCL-L-END | Leica | QBEND/10 |
| | CD248 | Purified | 564994 | BD | B1/35 |
| | CD45 | Purified | M0701 | Dako | 2B11 + PD7/26 |
| | CD3 | Purified | A0452 | Dako | Polyclonal Rabbit |
| | endomucin | Purified | sc-65495 | Santa cruz | (V.7C7) |

**Supplementary Table 2**. List of antibodies used for flow cytometry (FACS), Western Blot (WB) and immunohistochemistry (IHC) analysis.
