## Supplemental Table 3 for "Immune profiling of gliomas reveals a connection with Tau function and the tumor vasculature"

| Genes | Forward (5´-3´) | Reverse (3´-5´) |
|---|---|---|
| hP2RY12 | TGCCAAACTGGGACCAGGACCA | TGGTGGCTTCTGGTAGCGATC |
| hIBA1 | CCCTCCAAACTGGAAGGCTTCA | CTTTAGCTCAGGTGAGTCTTGG |
| hDLL3 | AAACCTATGGGCTTGAGGAG | CGCTGAGTACAATCAGTGGAA |
| hOLIG2 | CGGCTTTCCTCTATTTTGGTT | GTTACACGGCAGACGCTACA |
| hBCAM | CTGTCTGGAGAGAACTGCGT | TCTTGCAGGTGTAGGACAGG |
| hYKL40 | ACACCTGGGAGTGGAATGAT | AGTTCCATCCTCCGACAGAC |
| hCD44 | AGAAGGTGTGGGCAGAAGAA | AAATGCACCATTTCCTGAGA |
| hSERPINE1 | CATAGTGGAAGTGATAGAT | ACTCTGTTAATTCGTCTT |
| hCD34 | CCCTCAGTGTCTACTGCTGGTCT | GGAATAGCTCTGGTGGCTTGCA |
| hCD248 | AGACCACCACTCATTTGCCTGGAA | AGTTGGGATAATGGGAAGCGTGGT |
| hEMCN | GCAAGCACTTCAGCAACCAGCC | GGATCTGCCTTCCAGCACATTC |
| hIDH1 | CTATGATGGTGACGTTGCAGTCG | CCTCTGCTTCTACTGTCTTGCC |
| mEMCN | GCACACACCATGTCACTGCTTC | CAGCGCGATAACCACAGGCAAA |
| mCD248 | TTGATGGCACCTGGACAGAGGA | TCCAGGTGCAATCTCTGAGGCT |
| mCD3 | TCTCATTGCGGGACAGGATGGA | CCTTGGAGATGGCTGTACTGGT |

**Supplementary Table 3.** List of primers used for qRT-PCR analysis.
